# Nucleotide Analogues as Inhibitors of SARS-CoV-2 Polymerase

**DOI:** 10.1101/2020.03.18.997585

**Authors:** Minchen Chien, Thomas K. Anderson, Steffen Jockusch, Chuanjuan Tao, Shiv Kumar, Xiaoxu Li, James J. Russo, Robert N. Kirchdoerfer, Jingyue Ju

## Abstract

SARS-CoV-2, a member of the coronavirus family, is responsible for the current COVID-19 pandemic. Based on our analysis of hepatitis C virus and coronavirus replication, and the molecular structures and activities of viral inhibitors, we previously demonstrated that three nucleotide analogues inhibit the SARS-CoV RNA-dependent RNA polymerase (RdRp). Here, using polymerase extension experiments, we have demonstrated that the active triphosphate form of Sofosbuvir (a key component of the FDA approved hepatitis C drug EPCLUSA), is incorporated by SARS-CoV-2 RdRp, and blocks further incorporation. Using the same molecular insight, we selected the active triphosphate forms of three other anti-viral agents, Alovudine, AZT (an FDA approved HIV/AIDS drug) and Tenofovir alafenamide (TAF, an FDA approved drug for HIV and hepatitis B) for evaluation as inhibitors of SARS-CoV-2 RdRp. We demonstrated the ability of these three viral polymerase inhibitors, 3’-fluoro-3’-deoxythymidine triphosphate, 3’-azido-3’-deoxythymidine triphosphate and Tenofovir diphosphate (the active triphosphate forms of Alovudine, AZT and TAF, respectively) to be incorporated by SARS-CoV-2 RdRp, where they also terminate further polymerase extension. These results offer a strong molecular basis for these nucleotide analogues to be evaluated as potential therapeutics for COVID-19.

## Introduction

The COVID-19 pandemic, caused by SARS-CoV-2, has now spread to more than 100 countries on 6 continents. SARS-CoV-2 is a new member of the subgenus *Sarbecovirus*, in the Orthocoronavirinae subfamily, and is distinct from MERS-CoV and SARS-CoV.^1^ The coronaviruses are single-strand RNA viruses, sharing properties with other single-stranded RNA viruses such as hepatitis C virus (HCV), West Nile virus, Marburg virus, HIV virus, Ebola virus, dengue virus, and rhinoviruses. Coronaviruses, like HCV and the flaviviruses, are positive-sense single-strand RNA viruses,^2,3^ and these viruses share a similar replication mechanism requiring a RNA-dependent RNA polymerase (RdRp).

Potential inhibitors have been designed to target nearly every stage of the viral replication cycle.^2^ However, despite decades of research, no effective drug is currently approved to treat serious coronavirus infections such as SARS, MERS, and now COVID-19. One of the most important druggable targets for coronaviruses is the RdRp. This polymerase displays similar catalytic mechanisms and some key conserved amino acids in the active site among different positive sense RNA viruses, to which coronaviruses and HCV belong.^4^ Like RdRps in other viruses, the coronavirus enzyme is highly error-prone,^5^ which might increase its ability to accept modified nucleotide analogues as substrates. Nucleotide and nucleoside analogues that inhibit polymerases are an important group of anti-viral agents.^6–9^

Based on our analysis of hepatitis C virus and coronavirus replication, and the molecular structures and activities of viral inhibitors, we previously demonstrated that three nucleotide analogues inhibit the SARS-CoV RNA-dependent RNA polymerase (RdRp).^10^ Using polymerase extension experiments, we demonstrated that the activated triphosphate form of Sofosbuvir was incorporated by SARS-CoV RdRp, and blocked further incorporation. Using the same molecular insight, we selected two other anti-viral agents, Alovudine and AZT (the first FDA approved HIV/AIDS drug) for evaluation as inhibitors of SARS-CoV RdRp. Alovudine and AZT share a similar backbone structure (base and ribose) to Sofosbuvir, but have fewer modification sites and less steric hindrance. Furthermore, because these modifications on Alovudine and AZT are on the 3’ carbon in place of the OH group, they directly prevent further incorporation of nucleotides leading to permanent termination of RNA synthesis and replication of the virus. We also demonstrated the ability of two HIV reverse transcriptase inhibitors, 3’-fluoro-3’-deoxythymidine triphosphate and 3’-azido-3’-deoxythymidine triphosphate (the active triphosphate forms of Alovudine and AZT), to be incorporated by SARS-CoV RdRp where they also terminate further polymerase extension.^10^

In this paper, we first constructed SARS-CoV-2 RdRp based on a similar procedure to that of SARS-CoV,^11,12^ and then demonstrated that the above three nucleotide analogues (Fig. 1 a, b, d) are inhibitors of SARS-CoV-2 RdRp. Using structure-activity based molecular insight, we selected the active triphosphate form of Tenofovir alafenamide (TAF, Vemlidy, an acyclic adenosine nucleotide) (Fig. 1 c), which is an FDA approved drug for HIV and hepatitis B virus (HBV) infection, for evaluation as a SARS-CoV-2 RdRp inhibitor. The results indicated that the active triphosphate form of this molecule, tenofovir diphosphate (TFV-DP), also inhibited this polymerase. TAF, a prodrug form of the nucleotide analogue viral polymerase inhibitor, shows potent activity for HIV and HBV, but only limited inhibition of host nuclear and mitochondrial polymerases.^13,14^ It is activated by a series of hydrolases to the deprotected monophosphate form, TFV, and then by two consecutive kinase reactions to TFV-DP.^15^ TFV-DP is an acyclic nucleotide and does not have a 3’-OH group. Remarkably, this molecule is incorporated by both HIV and HBV polymerases, terminating nucleic acid elongation and viral replication.^13,15^ In addition, resistance mutations were rarely seen in patients treated with regimens including TAF.^16^ In view of the fact that the active triphosphate form of TAF, Tenofovir diphosphate, is much smaller than natural nucleoside triphosphates, we expect that it can easily fit within the active site of SARS-CoV-2 RdRp. As a noncyclic nucleotide, TFV-DP lacks a normal sugar ring configuration, and thus we reasoned that it is unlikely to be recognized by 3’-exonucleases involved in SARS-CoV-2 proofreading processes, decreasing the likelihood of developing resistance to the drug.^17^

**Fig. 1.**
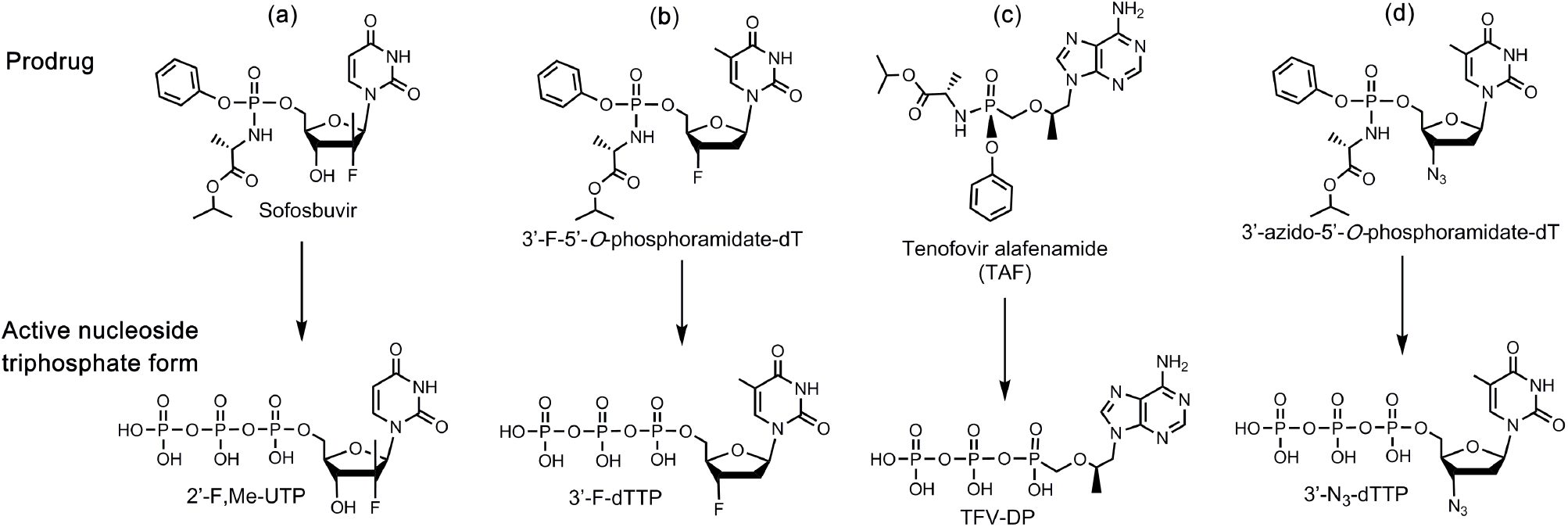
Structures of 4 prodrug viral inhibitors. Top: Prodrug (phosphoramidate) form; Bottom: Active triphosphate form.

## Results and Discussion

Given the 98% amino acid similarity of the SARS-CoV and SARS-CoV-2 RdRps,^10^ we reasoned that the four nucleotide analogues listed in Fig. 1 would also inhibit the SARS-CoV-2 polymerase. We thus assessed the ability of 2’-F,Me-UTP, 3’-F-dTTP, TFV-DP, and 3’-N_3_-dTTP (the active triphosphate forms of Sofosbuvir, Alovudine, TAF and AZT, respectively), to be incorporated by SARS-CoV-2 RdRp into an RNA primer and terminate the polymerase reaction.

The RdRp of SARS-CoV-2, referred to as nsp12, and its two protein cofactors, nsp7 and nsp8, whose homologs were shown to be required for the processive polymerase activity of nsp12 in SARS-CoV,^11,12^ were cloned and purified as described in the Methods. These three viral gene products in SARS-CoV-2 have high homology (e.g., 96% identity and 98% similarity for nsp12, with similar homology levels at the amino acid level for nsp7 and nsp8) to the equivalent gene products from SARS-CoV, the causative agent of SARS.^10^

We performed polymerase extension assays with 2’-F,Me-UTP, 3’-F-dTTP, 3’-N_3_-dTTP or TFV-DP + UTP, following the addition of a pre-annealed RNA template and primer to a pre-assembled mixture of the SARS-CoV-2 RdRp (nsp12) and two cofactor proteins (nsp7 and nsp8). The extended primer products from the reaction were subjected to MALDI-TOF-MS analysis. The RNA template and primer, corresponding to the 3’ end of the SARS-CoV-2 genome, were used for the polymerase assay, and their sequences are indicated at the top of Fig. 2. Because there are two A’s in a row in the next available positions of the template for RNA polymerase extension downstream of the priming site, if the 2’-F,Me-UTP, 3’-F-dTTP or 3’-N_3_-dTTP are incorporated by the viral RdRp, a single nucleotide analogue will be added to the 3’-end of the primer strand. If they are indeed inhibitors of the polymerase, the extension should stop after this incorporation; further 3’-extension should be prevented. Because the two A’s in the template are followed by 4 U’s, in the case of the TFV-DP/UTP mixture, two UTP’s should be incorporated prior to the incorporation and termination by TFV-DP, which has an adenosine base. As shown in Fig. 2, this is exactly what we observed. In the MALDI-TOF MS trace in Fig. 2a, a peak indicative of the molecular weight of a single base primer extension product with one 2’-F,Me-UTP was obtained (6644 Da observed, 6634 Da expected). Similarly, in the trace in Fig. 2b, a single extension peak indicative of a single base extension by 3’-F-dTTP is revealed (6623 Da observed, 6618 Da expected), with no further incorporation. In both of the above cases, the primer was nearly completely extended. In the trace in Fig. 2d, a single extension peak indicative of a single-base extension by 3’-N_3_-dTTP is seen (6633 Da observed, 6641 Da expected), with no evidence of further incorporation, though the incorporation efficiency was lower than for 2’-F,Me-UTP and 3’-F-dTTP, which may require further optimization. Finally, in the trace in Fig. 2c, a peak indicative of the molecular weight of a primer extension product formed by incorporating 2 U’s and 1 TFV is found (7198 Da observed, 7193 Da expected), in addition to other peaks representing partial incorporation (1 U, 6623 Da observed, 6618 Da expected) or misincorporation (3 U’s, 7235 Da observed, 7230 Da expected). Importantly, once the TFV-DP was incorporated, there was no further extension, indicating it was a permanent terminator.

**Fig. 2.**
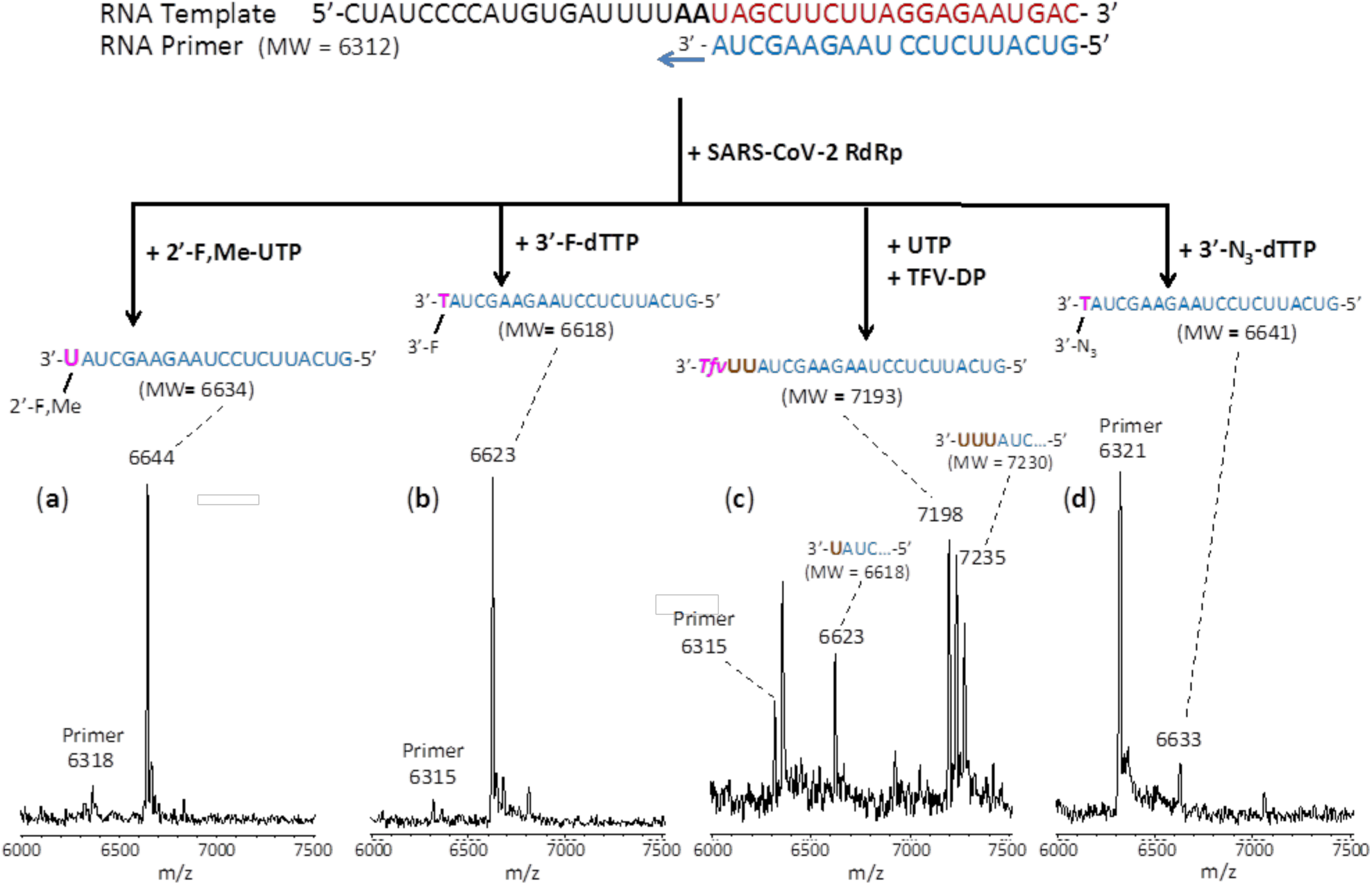
Incorporation of 2’-F,Me-UTP, 3’-F-dTTP, TFV-DP and 3’-N_3_-dTTP by SARS-CoV-2 RdRp to terminate the polymerase reaction. The sequences of the primer and template used for these extension reactions, which are at the 3’ end of the SARS-CoV-2 genome, are shown at the top of the figure. Polymerase extension reactions were performed by incubating (**a**) 2’-F,Me-UTP, (**b**) 3’-F-dTTP, (**c**) UTP + TFV-DP, and (**d**) 3’-N_3_-dTTP with pre-assembled SARS-CoV-2 polymerase (nsp12, nsp7 and nsp8), the indicated RNA template and primer, and the appropriate reaction buffer, followed by detection of reaction products by MALDI-TOF MS. The detailed procedure is shown in the Methods section. The accuracy for m/z determination is ± 10 Da.

In summary, these results demonstrate that the nucleotide analogues 2’-F,Me-UTP, 3’-F-dTTP, 3’-N_3_-dTTP, and TFV-DP are permanent terminators for the SARS-CoV-2 RdRp. Their prodrug versions (Sofosbuvir, 3’-F-5’-*O*-phosphoramidate dT nucleoside, 3’-N_3_-5’-*O*-phosphoramidate dT nucleoside, and TAF) are available or can be readily synthesized using the ProTide prodrug approach,^18^ as shown in Fig. 1 a-d, and can be developed as therapeutics for COVID-19. Importantly, since Sofosbuvir, Tenofovir and AZT are widely available FDA approved drugs, they can be evaluated quickly in laboratory and clinical settings for COVID-19 treatment.

## Methods

### Recombinant Protein Expression of RdRp (nsp12) and cofactors (nsp7 and nsp8) from SARS-CoV-2

#### SARS-CoV-2 nsp12

The SARS-CoV-2 sp12 gene was codon optimized and cloned into pFastBac with C-terminal additions of a TEV site and strep tag (Genscript). The pFastBac plasmid and DH10Bac *E. coli* (Life Technologies) were used to create recombinant bacmids. The bacmid was transfected into Sf9 cells (Expression Systems) with Cellfectin II (Life Technologies) to generate recombinant baculovirus. The baculovirus was amplified through two passages in Sf9 cells, and then used to infect 1 L of Sf21 cells (Expression Systems) and incubated for 48 hrs at 27°C. Cells were harvested by centrifugation, resuspended in wash buffer (25 mM HEPES pH 7.4, 300 mM NaCl, 1 mM MgCl_2_, 5 mM DTT) with 143 *μ*L of BioLock per liter of culture. Cells were lysed via microfluidization (Microfluidics). Lysates were cleared by centrifugation and filtration. The protein was purified using Strep Tactin superflow agarose (IBA). Strep Tactin eluted protein was further purified by size exclusion chromatography using a Superdex 200 Increase 10/300 column (GE Life Sciences) in 25 mM HEPES, 300 mM NaCl, 100 *μ*M MgCl_2_, 2 mM TCEP, at pH 7.4. Pure protein was concentrated by ultrafiltration prior to flash freezing in liquid nitrogen.

#### SARS-CoV-2 nsp7 and nsp8

The SARS-CoV-2 nsp7 and nsp8 genes were codon optimized and cloned into pET46 (Novagen) with an N-terminal 6x histidine tag, an enterokinase site, and a TEV protease site. Rosetta2 pLys *E. coli* cells (Novagen) were used for bacterial expression. After induction with isopropyl β-D-1-thiogalactopyranoside (IPTG), cultures were grown at 16°C for 16 hrs. Cells were harvested by centrifugation and pellets were resuspended in wash buffer (10mM Tris pH 8.0, 300 mM NaCl, 30 mM imidazole, 2 mM DTT). Cells were lysed via microfluidization and lysates were cleared by centrifugation and filtration. Proteins were purified using Ni-NTA agarose beads and eluted with wash buffer containing 300 mM imidazole. Eluted proteins were further purified by size exclusion chromatography using a Superdex 200 Increase 10/300 column (GE Life Sciences). Purified proteins were concentrated by ultrafiltration prior to flash freezing with liquid nitrogen.

### Extension reactions with RNA-dependent RNA polymerase

Oligonucleotides were purchased from IDT, Inc. The primer and template (sequences shown in Fig. 2) were annealed by heating to 70°C for 10 min and cooling to room temperature in 1x reaction buffer. The RNA polymerase mixture consisting of 6 *μ*M nsp12 and 18 *μ*M each of cofactors nsp7 and nsp8 was incubated for 15 min at room temperature in a 1:3:3 ratio in 1x reaction buffer. Then 5 *μ*l of the annealed template primer solution containing 2 *μ*M template and 1.7 *μ*M primer in 1x reaction buffer was added to 10 *μ*l of the RNA polymerase mixture and incubated for an additional 10 min at room temperature. Finally 5 *μ*l of a solution containing either 2 mM 2’-F,Me-UTP (a), 2 mM 3’-F-dTTP (b), 2 mM TFV-DP and 20 *μ*M UTP (c) or 3’-N_3_-dTTP (d) in 1x reaction buffer was added, and incubation was carried out for 2 hrs at 30°C. The final concentrations of reagents in the 20 *μ*l extension reactions were 3 *μ*M nsp12, 9 *μ*M nsp7, 9 *μ*M nsp8, 425 nM RNA primer, 500 nM RNA template, either 500 *μ*M 2’-F,Me-UTP (Sierra Bioresearch), 500 *μ*M 3’-F-dTTP (Amersham Life Sciences), 500 *μ*M TFV-DP/50 *μ*M UTP or 500 *μ*M 3’-N_3_-dTTP (Amersham Life Sciences), and 1x reaction buffer (10 mM Tris-HCl pH 8, 10 mM KCl, 2 mM MgCl_2_ and 1 mM β-mercaptoethanol). Following desalting using an Oligo Clean & Concentrator (Zymo Research), the samples were subjected to MALDI-TOF-MS (Bruker ultrafleXtreme) analysis.

## Acknowledgements

This research is supported by Columbia University and a grant from the Jack Ma Foundation to J.J. and the National Institute of Allergy and Infectious Disease AI123498 to R.N.K. A patent application on the work described has been filed.

## Author contributions

J.J. and R.N.K. conceived and directed the project; the approaches and assays were designed and conducted by J.J., X.L., S.K., S.J., J.J.R., M.C. and C.T., and SARS-CoV-2 polymerase and associated proteins nsp7 and 8 were cloned and purified by T.K.A. and R.N.K. Data were analyzed by all authors. All authors wrote and reviewed the manuscript.

## Additional information

**Correspondence** and requests for materials should be addressed to J.J. or R.N.K.

